## Supplement for "Evaluating the within-host dynamics of *Ranavirus* infection with mechanistic disease models and experimental data"

### I. Exploring Bayesian model diagnostics (for Model B2 only)

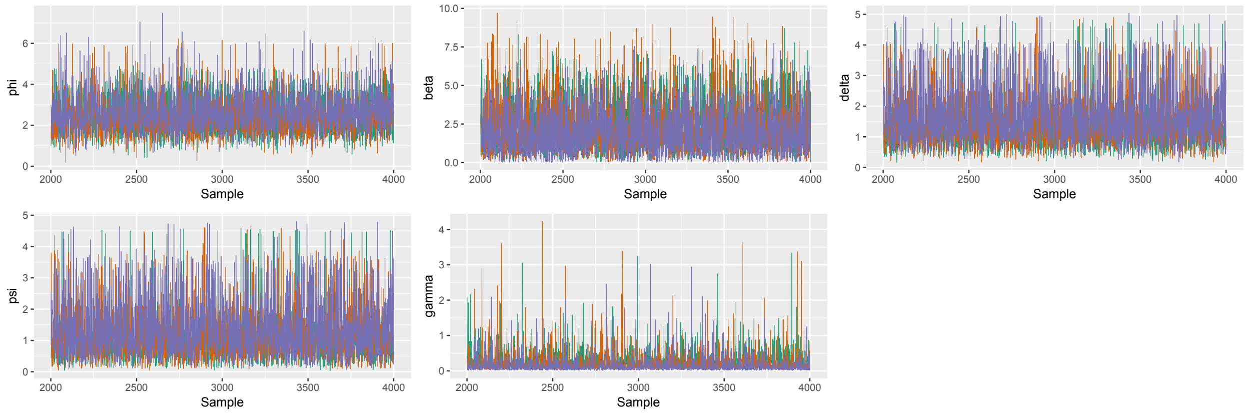


**Figure S1.** Traceplots of the HMC sampler for the five model parameters of model B2, with 3 sampling chains. All models converged after 4000 iterations.

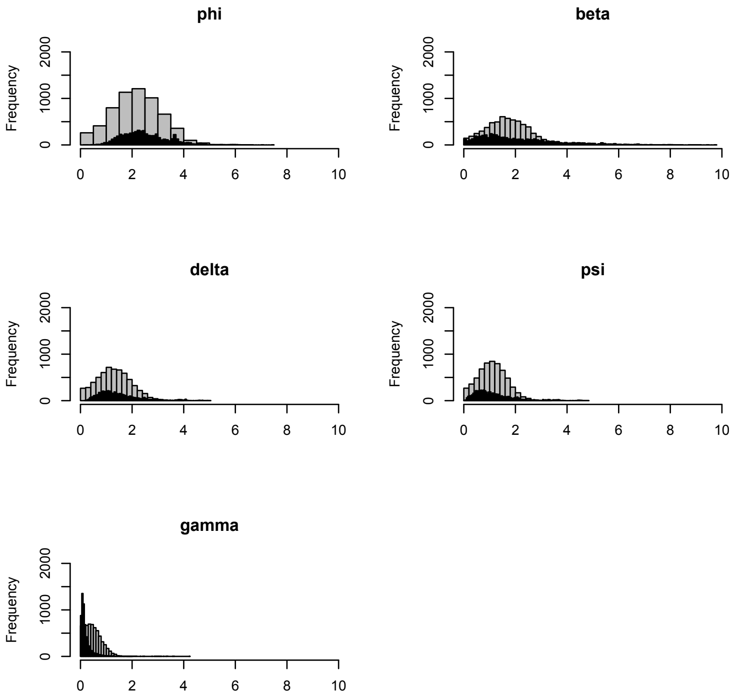


**Figure S2.** Comparison of prior probability distributions (gray bars) to marginal posterior samples for the five model parameters of model B2 (black bars). The bin size for the priors is larger to better illustrate the comparison. Also note that the model is estimating the means and standard deviations of a multivariate normal with the possibility of correlations among the parameters. We, however, detected no such correlations.

### II. Timing of Peak Titer

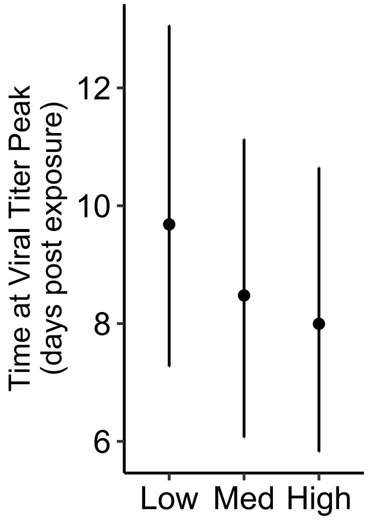


**Figure S3.** Effect of dosage on the predicted time of peak viral titer, derived from model B2. Median and 95% credible intervals are shown.

### III. Alternative model fits

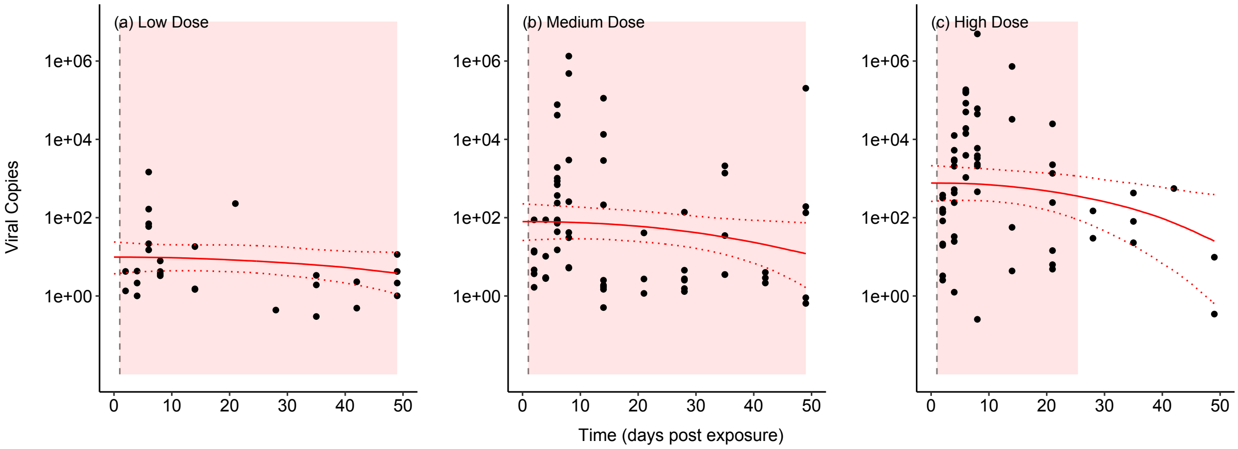


**Figure S4.** Fit of model A1 to the data with no dead individuals. Data and lines defined as in Figure 1 of the main text.

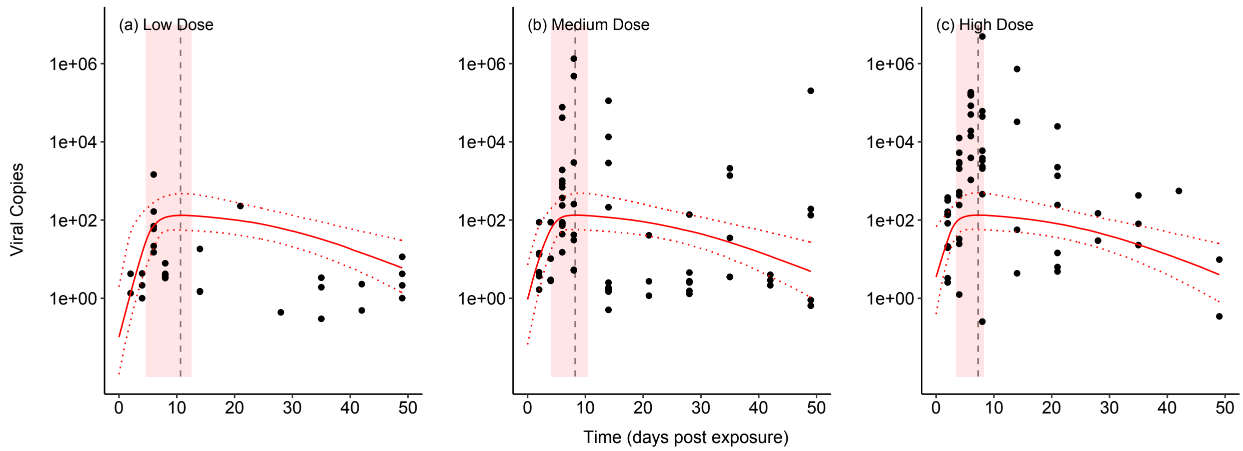


**Figure S5.** Fit of model A2 to the data with no dead individuals. This model fits poorly to the data from the high dose treatment, as it predicts the same peak titer for all three doses.

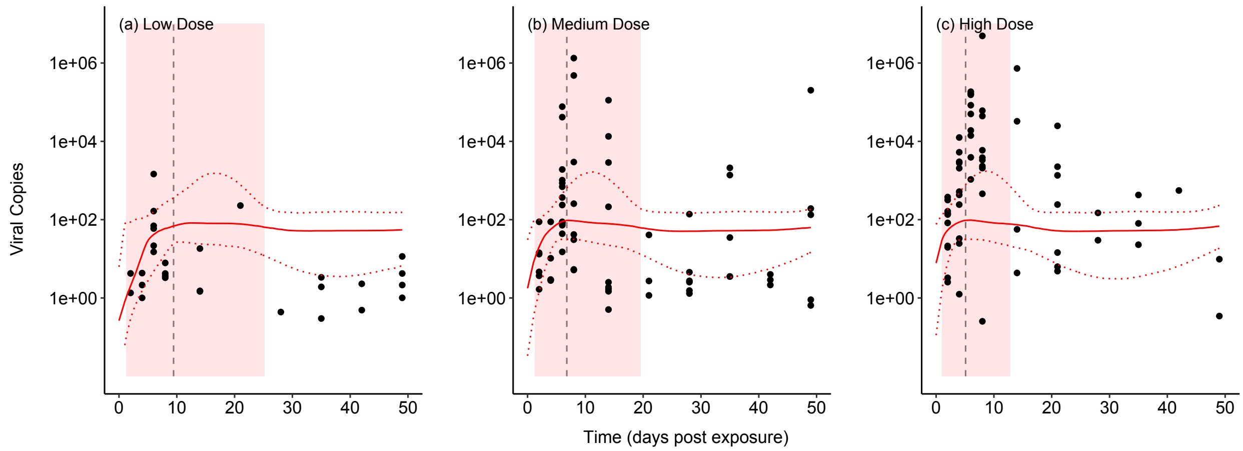


**Figure S6.** Fit of model B1 to the data with no dead individuals. This model struggled to converge, leading to multiple potential outcomes, as can be seen in the 95% credible interval of the model fit. This model also fits poorly to the high dose data, as it predicts the same peak titer for all doses.

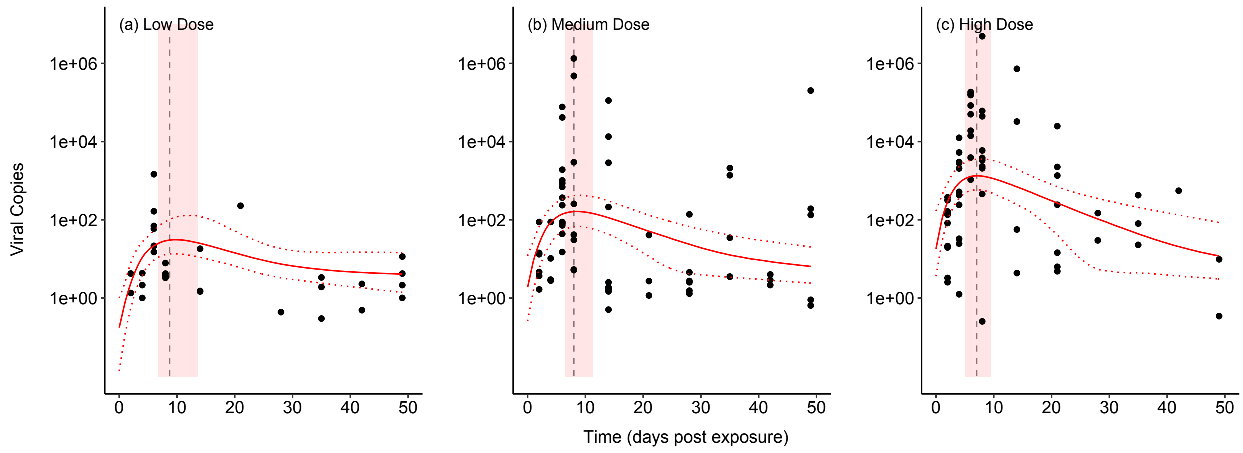


**Figure S7**. Fit of model B3 to the data with no dead individuals. This model is different from B2, because it assumes a logistic growth of the viral population (i.e., carrying capacity). However, it is clear that this model is over-fit, because it gives nearly identical predictions as model B2, but with an added parameter. Basically, the addition of the viral carrying capacity does not affect the predicted dynamics, and so the model is not parsimonious.

### IV. Model-fitting with dead individuals included

**Table S1.** Model structures and model comparisons for the data set that includes larvae that died of virus prior to their pre-determined sampling date. The bolded model (B2) is the most parsimonious based on LOO-IC selection.

| Class | ID | Structure | Penalty (*p*LOO) | LOO-IC | ΔLOO-IC |
|---|---|---|---|---|---|
| A | A1 | $V' = \phi V - \beta VZ$<br>$Z' = \psi Z \frac{V}{(V+\gamma)}$ | 8.6 | 1121.6 | 29.0 |
| | A2 | $V' = \phi V \left(1 - \frac{V}{K}\right) - \beta VZ$<br>$Z' = \psi Z \frac{V}{(V+\gamma)}$ | 7.6 | 1151.9 | 59.3 |
| B | B1 | $V' = \phi V - \beta VZ$<br>$Z' = (N_Z - \delta Z) + \psi ZV$ | 10.8 | 1144.5 | 51.9 |
| | **B2** | $V' = \phi V - \beta VZ$<br>$Z' = (N_Z - \delta Z) + \psi Z \frac{V}{(V+\gamma)}$ | **10.0** | **1092.6** | **0** |
| | B3 | $V' = \phi V \left(1 - \frac{V}{K}\right) - \beta VZ$<br>$Z' = (N_Z - \delta Z) + \psi Z \frac{V}{(V+\gamma)}$ | 10.0 | 1092.9 | 0.3 |

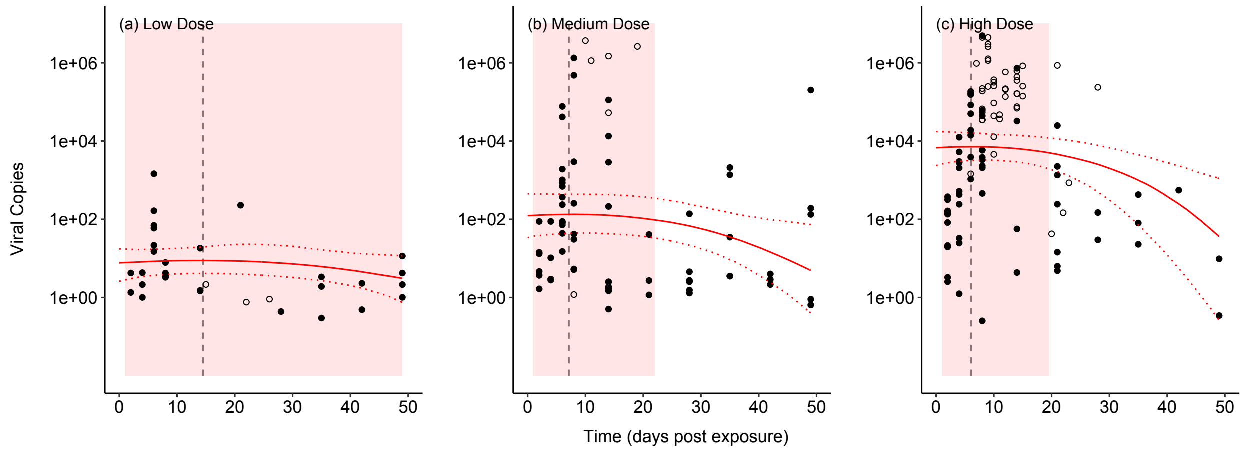


**Figure S8.** Fit of model A1 to the data. Data include individuals that died before their pre-determined sampling data (open circles), and lines are defined as in Figure 1 of the main text.

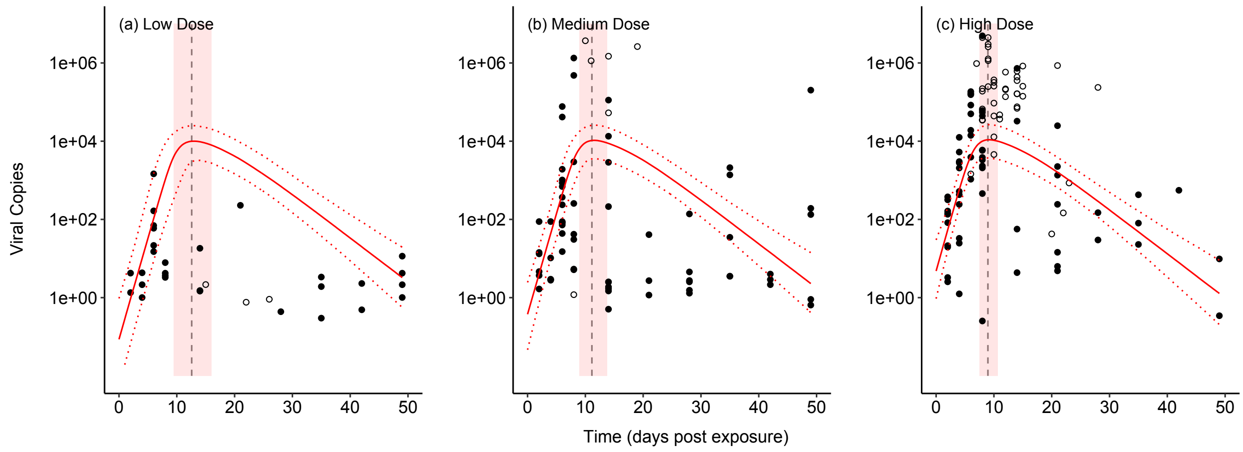


**Figure S9.** Fit of model A2 to the data. When dead individuals are included, this model fits poorly to the data from the low dose treatment. Also, it predicts the same peak titer for all three doses.

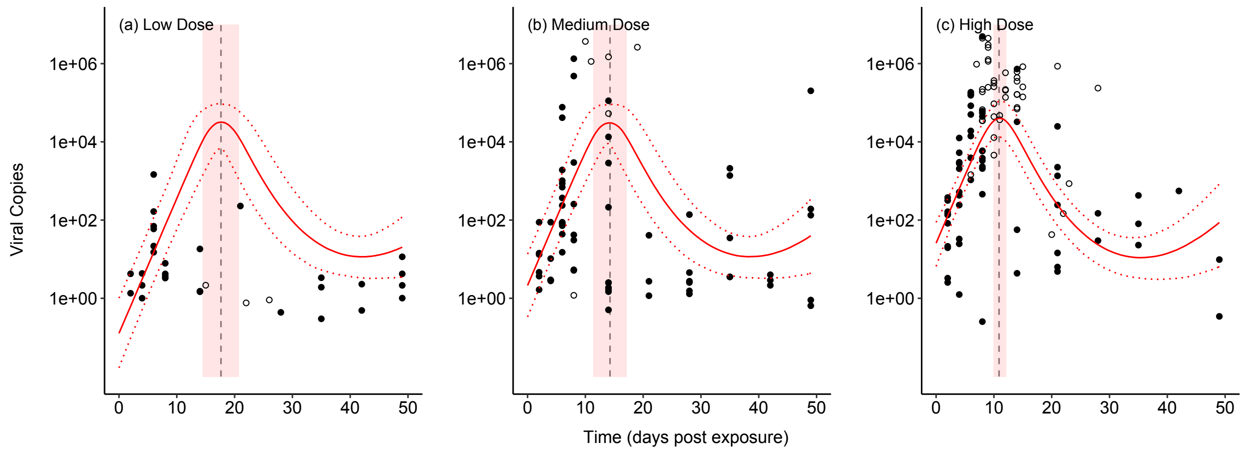


**Figure S10.** Fit of model B1 to the data. This model predicts the same peak titer for all three doses. It also predicts damped oscillations towards a stable equilibrium, which is why the predicted titer is rising at the end. This doesn't seem biologically realistic, but we don't have enough long-term data to fully test this prediction. In any case, the model does not have the best fit, because of its poor fit to the data from the low dose treatment.

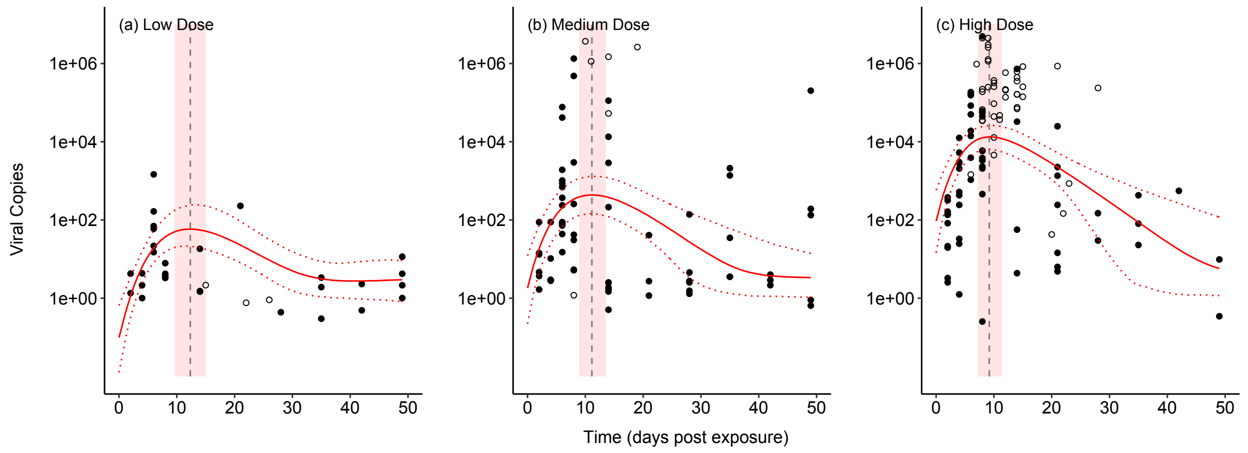


**Figure S11**. Fit of model B3 to the data. This model is different from B2, because it assumes a logistic growth of the viral population (i.e., carrying capacity). However, it is clear that this model is over-fit, because it gives nearly identical predictions as model B2, but with an added parameter. Basically, the addition of the viral carrying capacity does not affect the predicted dynamics, and so the model is not parsimonious.
